# Vaccination against the digestive enzyme Cathepsin B using a YS1646 *Salmonella enterica* Typhimurium vector provides almost complete protection against *Schistosoma mansoni* challenge in a mouse model

**DOI:** 10.1101/652644

**Authors:** Adam S Hassan, Nicholas H Zelt, Dilhan J Perera, Momar Ndao, Brian J Ward

**Author notes:** Co-corresponding authors (BJW) (MN).

## Abstract

*Schistosoma mansoni* threatens hundreds of millions of people in >50 countries. Schistosomulae migrate through the lung and adult worms reside adjacent to the intestinal mucosa. None of the candidate vaccines in current development is designed to elicit a mucosal response. We have repurposed an attenuated *Salmonella enterica* Typhimurium strain (YS1646) to produce such a vaccine targeting Cathepsin B (CatB), a digestive enzyme important for parasite survival. Promoter-Type 3 secretory signal pairs were screened for protein expression *in vitro* and transfected into YS1646 to generate candidate vaccine strains. Two strains were selected for *in vivo* evaluation (nirB_SspH1 and SspH1_SspH1). Female C57BL/6 mice were immunized twice, 3 weeks apart, using six strategies: i) saline gavage (control), ii) the ‘empty’ YS1646 vector orally (PO) followed by intramuscular recombinant CatB (20μg IM rCatB), iii) two doses of IM rCatB, iv) two PO doses of YS1646-CatB, v) IM rCatB then PO YS1646-CatB and vi) PO YS1646-CatB then IM rCatB. Serum IgG responses to CatB were monitored by ELISA. Three weeks after the second dose, mice were challenged with 150 cercariae and sacrificed 7 weeks later to assess adult worm and egg burden (liver and intestine), granuloma size and egg morphology. CatB-specific IgG antibodies were low/absent in the control and PO only groups but rose substantially in other groups (5898-6766ng/mL). The highest response was in animals that received nirB_SspH1 YS1646 PO then IM rCatB. In this group, reductions in worm and intestine/liver egg burden (vs. control) were 93.1% and 79.5%/90.3% respectively (all *P*<.0001). Granuloma size was reduced in all vaccinated groups (range 32.86–52.83 ×10^3^μm^2^) and most significantly in the nirB_SspH1 + CatB IM group (34.74±3.35 ×10^3^μm^2^vs. 62.22±6.08 ×10^3^μm^2^: vs. control *P*<.01). Many eggs in the vaccinated animals had abnormal morphology. Targeting CatB using a multi-modality approach can provide almost complete protection against *S. mansoni* challenge.

**Author Summary:** Schistosomiasis is a parasitic disease that affects over 250 million people worldwide and over 800 million are at risk of infection. Of the three main species, *Schistosoma mansoni* is the most widely distributed and is endemic in the Caribbean, South America, Africa, and the Middle East. It causes a chronic disease with severe negative effects on quality of life. Mass drug administration of praziquantel is the only available course of action due to a current lack of vaccines. However, praziquantel does not protect from reinfection. Therefore, a vaccine would be beneficial as a long-term solution to reduce morbidity and transmission of the disease. Our group has repurposed the attenuated YS1646 strain of *Salmonella* Typhimurium as an oral vaccine vector for the digestive enzyme Cathepsin B of *S. mansoni*. Oral vaccination followed by an intramuscular dose of recombinant Cathepsin B lead to significant reductions in parasite burden in mice. These animals had the highest titers in serum IgG and intestinal IgA antibodies. This multimodal vaccination approach also elicited both Th1 and Th2 cytokines as seen by the increases in IFNγ and IL-5. Finally, vaccinated mice had reductions in granuloma size along with a higher proportion of morphologically-abnormal eggs. This work demonstrates that a YS1646-based, multimodality, prime-boost immunization schedule can provide nearly complete protection against *S. mansoni* in a well-established murine model.

## Introduction

Schistosomiasis is caused by a number of *Schistosoma spp*. These trematodes currently infect >250 million people worldwide and more than 800 million are at risk of infection [1]. The World Health Organization (WHO) considers schistosomiasis to be the most important human helminth infection in terms of mortality and morbidity [2]. Of the three main human schistosome species, *S. mansoni* is the most widespread; causing a significant burden of disease in South America, Sub-Saharan Africa, the Caribbean and the Middle East [3].

The current treatment of schistosomiasis relies heavily on the drug praziquantel. This oral anthelminthic paralyzes the adult worms and has a reported efficacy of 85-90% [4]. The availability of only one effective drug is a precarious situation however and praziquantel resistance has been observed both experimentally [5, 6] and in the field [7, 8]. Furthermore, praziquantel treatment does not prevent re-infection. There is a clear need for a vaccine that can be used in conjunction with mass drug administration (MDA) and vector control efforts.

The WHO Special Program for Research and Training in Tropical Diseases (TDR/WHO) has encouraged the search for a vaccine that can provide ≥40% protection against *S. mansoni* [9]. Despite this relatively ‘low bar’, few candidate vaccines have achieved >50% protection in murine or other animal models [10] and even fewer have progressed to human trials [11]. Our group has previously demonstrated 60-70% protection in a *S. mansoni* murine challenge model by targeting Cathepsin B using intramuscular (IM)-adjuvanted formulations [12, 13]. Cathepsin B (CatB) is a cysteine protease found in the cecum of both the migratory larval form of *S. mansoni* (ie: the schistosomula) and in the gut of the adult worm. CatB is important for the digestion of host blood macromolecules such as hemoglobin, serum albumin and immunoglobulin G (IgG) [14]. Suppression of CatB expression using RNA interference (RNAi) has a major impact on parasite growth and fitness [15]. Because the schistosomulae migrate through the lungs and the adult worms reside adjacent to the gut mucosa, we wished to determine if a vaccination strategy that targeted induction of both mucosal and systemic responses to CatB could improve protection.

YS1646 is a highly attenuated *Salmonella enterica* serovar Typhimurium carrying mutations in the msbB (lipopolysaccharide or LPS) and purI (purine biosynthesis pathway) genes that was originally developed as a possible cancer therapeutic [16]. Although its development was halted when it failed to provide benefit in a large phase I trial in subjects with advanced cancer, it was well-tolerated when administered intravenously at doses of up to 3.0×10^8^ colony-forming units/m^2^ [16]. We are seeking to repurpose YS1646 as a novel vaccination platform and reasoned that a locally-invasive but highly attenuated *Salmonella* vector might induce both local and systemic responses to CatB. The flagellin protein of *S.* Typhimurium has been proposed as a general mucosal adjuvant through its action on toll-like receptor (TLR) 5 [17]. Other *Salmonella* products such as LPS would be expected to further enhance immune responses by triggering TLR4 [18, 19]. Indeed, live attenuated *Salmonella* have multiple potential advantages as vaccine vectors and have been used to express foreign antigens against infectious diseases and cancers [20–22]. They directly target the intestinal M cells overlying the gut-associated lymphoid tissues (GALT) [21, 23–26], have large ‘carrying’ capacity [27] and are easy to manipulate both in the laboratory and at industrial scale. Although there is considerable experience with the attenuated *S. typhi* vaccine strain (Ty21a: Vivotif™) in the delivery of heterologous antigens [21, 28], far less is known about the potential of other *Salmonella* strains. Of direct relevance to the current work, Chen and colleagues used YS1646 to express a chimeric *S. japonicum* antigen that induced both strong antibody and cellular responses after repeated oral dosing and provided up to 62% protection in a murine challenge model [29].

In this work, we exploited *Salmonella* type-III secretory signals (T3SS) and both T3SS-specific and constitutive promoters to generate a panel of YS1646 strains with plasmid-based expression of enhanced green fluorescent protein (eGFP) or full-length CatB. This panel was screened for protein expression in monomicrobial culture and murine RAW 264.7 murine macrophages and the most promising constructs were advanced to *in vivo* testing in adult female C57BL/6 mice. Animals were vaccinated with the two most promising strains using several strategies and then subjected to cercarial challenge. Herein we report that a two-dose, multimodality schedule starting with oral (PO) gavage of YS1646 bearing the nirB_SspH1_CatB plasmid followed by intramuscular (IM) recombinant CatB (rCatB) was able to reduce both worm and tissue egg burden by 80-90%. Such reductions are among the best ever reported for any *S. mansoni* candidate vaccine in a murine model.

## Methods

### Ethics statement

All animal procedures were conducted in accordance with Institutional Animal Care and Use Guidelines and were approved by the Animal Care and Use Committee at McGill University (Animal Use Protocol 7625). These guidelines adhere to those of the Canadian Council on Animal Care (CCAC).

### Plasmids

Gene segments of the *pagc* promoter as well as the *sopE2, sspH1, sspH2, sptP, steA, steB* and *steJ* promoters and secretory signals were cloned from YS1646 genomic DNA (American Type Culture Collection, Manassas, VA) and the *nirB* and *lac* promoters were cloned from *E. coli* genomic DNA (strain AR_0137) (ThermoFischer Scientific, Eugene, OR). *S. mansoni* CatB complementary DNA (cDNA) was sequence-optimized for expression in *S. enterica* Typhimurium [Java Codon Optimization Tool (jcat)], synthesized by GenScript (Piscataway, NJ) and inserted into the pUC57 plasmid with a 6x His tag at the 3’ end. Promoter-T3SS pairs were cloned upstream of the CatB gene and inserted separately into pQE30 (Qiagen, Hilden, Germany). Parallel constructs were made with CatB gene replaced by eGFP to produce expression plasmids used for imaging studies. See Figure 1 for the general plasmid map and Table 1 for a summary of the expression cassettes produced. All plasmids were sequenced to verify successful cloning (McGill Genome Centre, Montreal, QC). *S. enterica* Typhimurium YS1646 (Cedarlane Labs, Burlington, ON) was cultured in Lysogeny broth (LB) media and strains bearing each construct were generated by electroporation (5ms, 3kV: Biorad, Hercules, CA). Successfully transformed strains were identified using LB agar containing 50 μg/mL ampicillin (Wisent Bioproducts, St-Bruno, QC). Aliquots of each transformed strain were stored in LB with 15% glycerol at −80°C until used in experiments.

**Fig 1.**
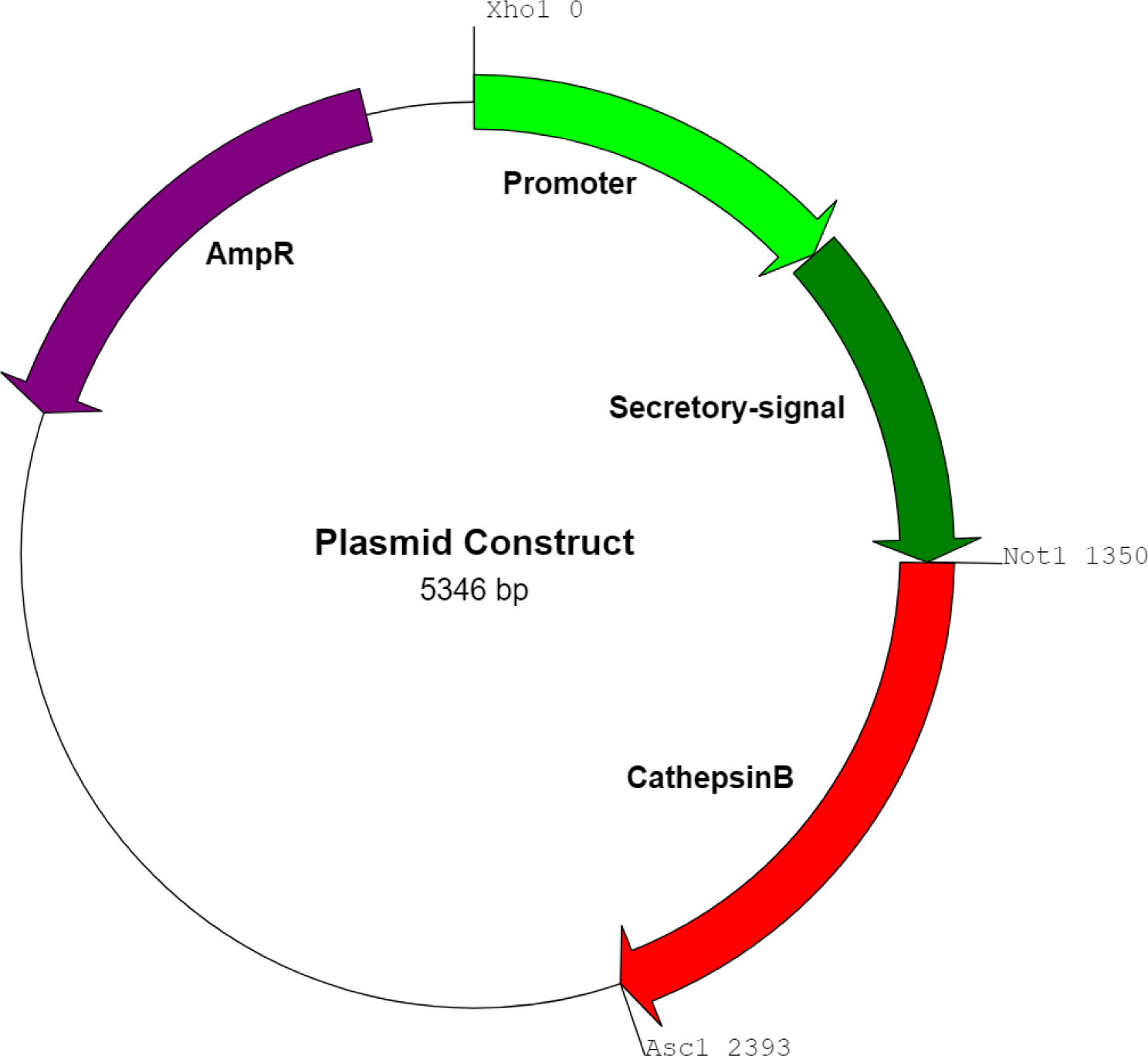
Plasmid map for recombinant YS1646 strains. The pQE-30 plasmid served as a backbone. The promoter and secretory signal were inserted between the Xho1 and Not1 restriction sites. The full-length Cathepsin B gene was inserted between the Not1 and Asc1 sites. An ampicillin resistance gene was used as a selectable marker.

**Table 1.**
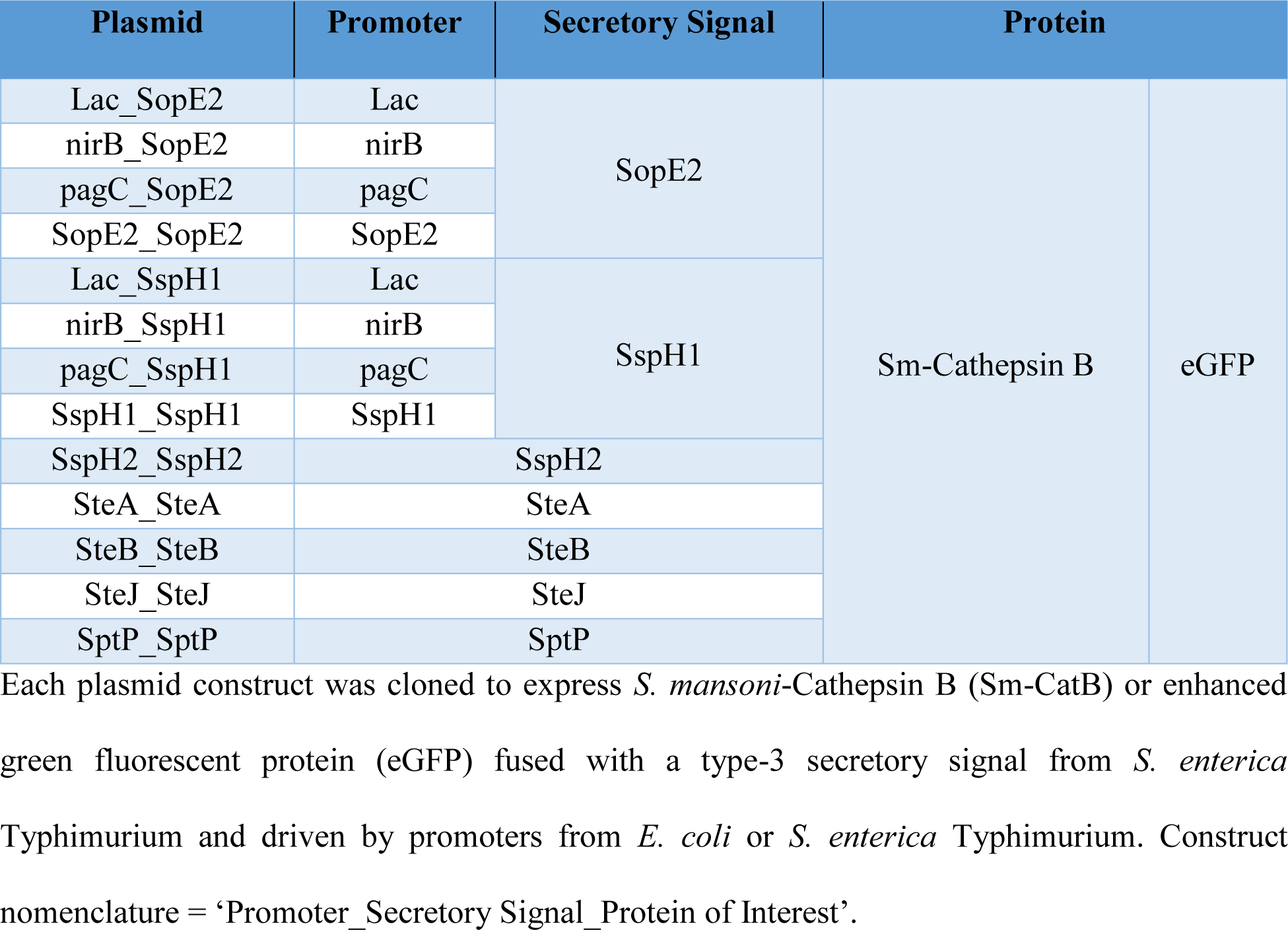
Recombinant *Salmonella* constructs.

### Western blotting

Recombinant YS1646 strains were grown in LB broth with 50 μg/mL ampicillin at 37°C in a shaking incubator under aerobic or low oxygen (sealed twist-cap tubes) conditions. Bacterial lysates were prepared by centrifugation (9,000xg for 5 min) then boiling the pellet (100°C × 10 min). Proteins from the culture supernatant were precipitated with 10% trichloroacetic acid for 1 hour on ice followed by centrifugation (9,000xg for 2 min) and removal of the supernatant. Protein pellets were resuspended in NuPAGE LDS sample buffer and NuPAGE reducing agent according to the manufacturer’s instructions (Thermo Fisher). Immunoblotting was performed as previously described [12]. Briefly, samples were run on a 4-12% Bis-Tris PAGE gel and transferred to nitrocellulose membranes (Thermo Fisher). Membranes were incubated in blocking buffer (5% skim milk in PBS [pH 7.4; 0.01M phosphate buffer, 0.14 M NaCl]) for 1 hour at room temperature (RT) with gentle agitation then washed three times in wash buffer (PBS [pH 7.4; 0.01M phosphate buffer, 0.14 M NaCl], 0.1% Tween 20 (Sigma-Aldrich, St. Louis, MO). Membranes were incubated with a murine, monoclonal anti-polyhistidine primary antibody (1:2,500; Sigma-Aldrich) in blocking buffer overnight at 4°C with gentle shaking. Membranes were washed three times in wash buffer then incubated with a goat, anti-mouse IgG-horseradish peroxidase secondary antibody (1:5000; Sigma-Aldrich) in blocking buffer for 1 hour at RT with gentle agitation. Membranes were washed three times followed by addition of Supersignal West Pico chemiluminescent substrate (Thermo Fisher) as per the manufacturer’s instructions and developed using an autoradiography cassette and the X-OMAT 2000 processor system (Kodak, Rochester, NY).

### *In vitro* macrophage infection

Murine macrophage-like cells (RAW 264.7: ATCC-TIB 71) were seeded at 10^6^ cells/well in 12-well plates in Dulbecco’s Modified Eagle’s medium (DMEM) (Wisent Bioproducts) supplemented with 10% fetal bovine serum (FBS: Wisent Bioproducts). Transformed YS1646 were diluted in DMEM-FBS to give a multiplicity of infection of 100 and centrifuged onto the monolayer (110xg for 10 min) to synchronize the infection. After 1 hour at 37°C in 5% CO_2_, plates were washed three times with phosphate buffered saline (PBS: Wisent Bioproducts) and replaced in the incubator with DMEM-FBS containing 50 μg/mL gentamicin (Sigma-Aldrich) to kill any extracellular bacteria and prevent re-infection. After 2 hours, the cells were washed with PBS three times and the gentamicin concentration was lowered to 5 μg/mL. After 24 hours, the cells were harvested, transferred to Eppendorf tubes and centrifuged (400xg for 5 min). Pellets were prepared for western blotting as above. For imaging experiments, RAW 264.7 cells were seeded into 6-well chamber slides at 10^4^ cells/well and cultured as above. After 24 hours, the cells were stained with 4’,6-diamidino-2-phenylindole (DAPI) (Thermo Fisher), fixed with 4% paraformaldehyde in PBS and incubated for 10 min at RT. Images were obtained using a Zeiss LSM780 laser scanning confocal microscope and analyzed using ZEN software (Zeiss, Oberkochen, Germany).

### Purification of recombinant cathepsin B

*S. mansoni* CatB was cloned and expressed in *Pichia pastoris* as previously described [12]. Briefly, the yeast cells were cultured at 28°C with shaking in buffered complex glycerol medium (BMGY) (Fisher Scientific, Ottawa, ON). After two days, cells were pelleted (3,000xg for 5 min) and resuspended in fresh BMMY to induce protein expression. After 3 further days of culture, cells were harvested (3,000xg for 5 min) and supernatants were collected and purified by Ni-NTA affinity chromatography. Immunoblotting for the His-tag (as above) confirmed successful expression of CatB. Protein concentration was estimated by Piece bicinchoninic acid assay (BCA) (Thermo Fisher) and aliquots of the rCatB were stored at −80°C until used.

### Immunization protocol

Female 6–8 week old C57BL/6 mice were purchased from Charles River Laboratories (Senneville, QC). All animals received two doses three weeks apart (See Fig 2 for experimental design). Oral dosing (PO) was accomplished by gavage three times every other day (200 μL containing 1×10^9^ colony-forming units (CFUs)/dose). Intramuscular (IM) vaccinations were administered using a 25g needle in the lateral thigh (20μg rCatB in 50μL PBS). Each experiment included six groups with 8 mice/group: i) saline PO twice (Control or PBS) ii) YS1646 transformed with ‘empty’ pQE30 vector (EV) PO followed by rCatB IM (EV→IM), iii) CatB-bearing YS1646 PO twice (PO→PO), iv) rCatB IM twice (IM→IM), v) CatB-bearing YS1646 PO followed by rCatB IM (PO→IM), and vi) rCatB IM followed by CatB-bearing YS1646 PO (IM→PO).

**Figure 2.**
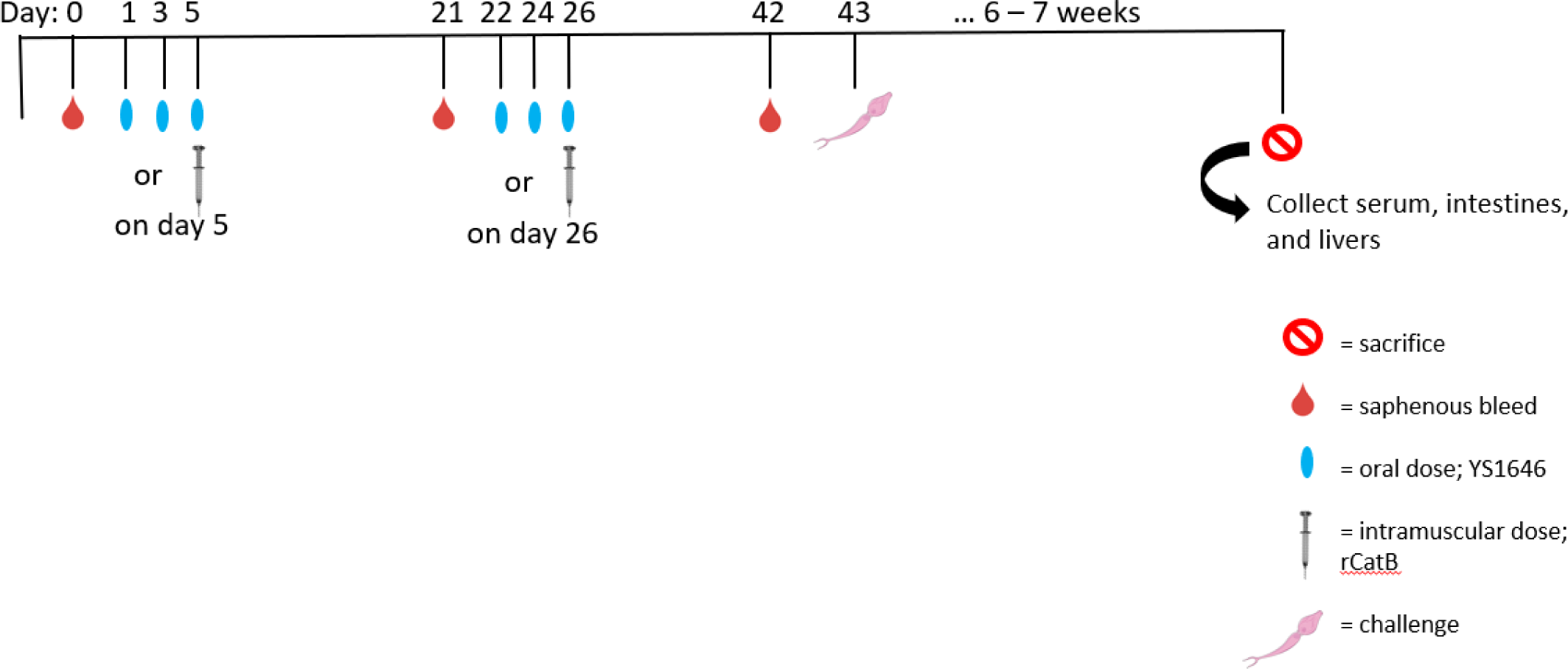
Immunization schedule. Baseline serum was collected on day 0 for all mice. Depending on the experimental group, mice receive 3 oral doses of YS1646 (1×10^9^ cfu/dose) or PBS every other day while others receive an intramuscular dose of 20 μg of CatB on day 5. Mice were bled and underwent a second round of vaccination three weeks later before being challenged with 150 *S. mansoni* cercariae by tail penetration. All animals were sacrificed 6 – 7 weeks post-infection.

### Intestine processing for IgA assessment

Four weeks after the second vaccination, the animals were sacrificed, and 10 cm of the proximal small intestine was collected. Tissue was weighed and stored in a protease inhibitor cocktail (Sigma Aldrich) at a 1:5 dilution (w/v) on ice until processed. Tissue was homogenized (Homogenizer 150; Fisher Scientific), centrifuged at 2500xg at 4°C for 30 minutes and the supernatant was collected. Supernatants were stored at −80°C until analyzed by ELISA.

### Humoral response by enzyme-linked immunosorbent assay (ELISA)

#### Serum IgG and intestinal IgA

Blood was collected from the saphenous vein at baseline (week 0) and at 3 and 6 weeks in microtainer serum separator tubes (BD Biosciences, Mississauga, ON, Canada). Cleared serum samples were obtained following the manufacturer’s protocol and stored at −20°C until used. Serum CatB-specific IgG and intestinal CatB-specific IgA levels were assessed by ELISA as previously described [30]. Briefly, U-bottom, high-binding 96-well plates (Greiner Bio-One, Frickenhausen, Germany) were coated overnight at 4°C with rCatB (0.5 μg/mL) in 100 mM bicarbonate/carbonate buffer at pH 9.6 (50 μL/well). Each plate contained a standard curve with 2-fold dilutions of purified mouse IgG or IgA (Sigma Aldrich, St. Louis, MO) starting at 2,000 ng/mL. The plates were washed three times with PBS (pH 7.4) and incubated with blocking buffer (2% bovine serum albumin (Sigma-Aldrich) in PBS-Tween 20 (0.05%; Fisher Scientific)) at 37°C for 1 hour. The plates were washed three times with PBS and diluted serum samples (1:50 in blocking buffer) were added in duplicate (50 μL/well). Blocking buffer was added to the standard curve wells. After 1 hour at 37°C, the plates were washed with PBS four times and horseradish peroxidase-conjugated anti-mouse IgG or horseradish peroxidase-conjugated anti-mouse IgA (Sigma Aldrich) diluted 1: 20,000 (1:10,000 for IgA) in blocking buffer was added for 30 min (IgG) or 1 hour (IgA) at 37°C (75 μL/well). Plates were washed with PBS six times and 3,3′,5,5′-Tetramethyl benzidine (TMB) substrate (100 μL/well; Millipore, Billerica, MA) was used for detection followed by 0.5 M H_2_SO_4_ after 15 min (50 μl/well; Fisher Scientific). Optical density (OD) was measured at 450 nm with an EL800 microplate reader (BioTek Instruments Inc., Winooski, VT). The concentration of CatB-specific IgG and IgA were calculated by extrapolation from the mouse IgG or IgA standard curves.

#### Serum IgG1 and IgG2c

Serum CatB-specific IgG1 and IgG2c levels were assessed by ELISA as previously described [12]. Briefly, Immulon 2HB flat-bottom 96-well plates (Thermo Fisher) were coated overnight at 4°C with rCatB (0.5 μg/mL) in 100 mM bicarbonate/carbonate buffer at pH 9.6 (50 μL/well). The plates were washed three times with PBS-Tween 20 (PBS-T: 0.05%; Fisher Scientific) and were blocked as above for 90 min. Serial serum dilutions in duplicate were incubated in the plates for 2 hours. Control (blank) wells were loaded with PBS-T. After washing three times with PBS-T, goat anti-mouse IgG1-horseradish peroxidase (HRP) (Southern Biotechnologies Associates, Birmingham, AL) and goat anti-mouse IgG2c-HRP (Southern Biotechnologies Associates) were added to the plates and incubated for 1 hour at 37°C. After a final washing step, TMB substrate (50 μL/well; Millipore, Billerica, MA) was used for detection followed by 0.5 M H_2_SO_4_ after 15 min (25 μl/well; Fisher Scientific). Optical density (OD) was measured at 450 nm with an EL800 microplate reader (BioTek Instruments Inc.). The results are expressed as the mean IgG1/IgG2c ratio of the endpoint titers ± standard error of the mean. Endpoint titers refer to the reciprocal of the highest dilution that gives a reading above the cut-off calculated as previously described [31].

### Cytokine production by multiplex ELISA

In some experiments, some of the animals were sacrificed 4 weeks after the second vaccination. Spleens were collected and splenocytes were isolated as previously described with the following modifications [13]. Splenocytes were resuspended in 96-well plates (10^6^ cells/well) in RPMI-1640 (Wisent Bioproducts) supplemented with 10% fetal bovine serum, 1 mM penicillin/streptomycin, 10 mM HEPES, 1X MEM non-essential amino acids, 1 mM sodium pyruvate, 1 mM L-glutamine (all from Wisent Bioproducts), 0.05 mM 2-mercaptoethanol (Sigma-Aldrich). The cells were incubated at 37°C in the presence of 2.5 μg/mL of rCatB for 72 hours after which the supernatant cytokine levels were measured by QUANSYS multiplex ELISA (9-plex) (Quansys Biosciences, Logan, UT) following the manufacturer’s recommendations.

### *Schistosoma mansoni* challenge

*Biomphalaria glabrata* snails infected with the *S. mansoni* Puerto Rican strain were obtained from the Schistosomiasis Resource Center of the Biomedical Research Institute (Rockville, MD) through NIH-NIAID Contract HHSN272201700014I for distribution through BEI Resources. Mice were challenged three weeks after the second immunization (week 6) with 150 cercariae by tail exposure and were sacrificed seven weeks post-challenge as previously described [32]. Briefly, adult worms were counted after perfusion of the hepatic portal system and manual removal from the mesenteric veins. The livers and intestines were harvested from each mouse, weighed and digested in 4% potassium hydroxide overnight at 37°C. The next day, the number of eggs per gram of tissue was recorded by microscopy. A small portion of each liver was placed in 10% buffered formalin phosphate (Fisher Scientific) and processed for histopathology to assess mean granuloma size and egg morphology (H&E staining). Granuloma area was measured using Zen blue software (version 2.5.75.0; Zeiss). Briefly, working at 400x magnification, 6 – 8 granulomas per mouse with a clearly visible egg were traced using the screen stylus and mean areas were presented as x10^3^ μm^2^ ± SEM. Eggs were classified as abnormal if obvious shrinkage had occurred, if internal structure was lost or if the perimeter of the egg was crenelated and are reported as a percent of the total eggs counted (± SEM).

### Statistical analysis

Statistical analysis was performed using GraphPad Prism 6 software (La Jolla, CA). In each experiment, reductions in worm and egg burden were expressed relative to the saline control group numbers. Data were analyzed by one-way ANOVA. P values less than 0.05 were considered significant.

## Results

### In vitro Expression and Secretion of CatB by Transformed YS1646 Strains

Thirteen expression cassettes were built and the sequences were verified (McGill University Genome Quebec Innovation Centre) (Table 1). The promoter/T3SS pairs were inserted in-frame with either *S. mansoni* CatB or eGFP. In monomicrobial culture, CatB expression was effectively driven by the nirB_SspH1, SspH1_SspH1 and SteA_SteA plasmids (Fig 3A) with the greatest production from the nirB promoter in low oxygen conditions as previously reported [29]. Secreted CatB was detectable in the monomicrobial culture supernatants only with YS1646 bearing the SspH1_SspH1 construct (Fig 3A). In infected RAW 264.7 cells, all of the constructs produced detectable eGFP by immunofluorescence (Fig 3B) but only the YS1646 bearing the nirB_SspH1 and SspH1_SspH1 constructs produced CatB detectable by immunoblot (Fig 3C). These constructs also led to the greatest eGFP expression in the RAW 264.7 cells and so were selected for *in vivo* testing.

**Figure 3.**
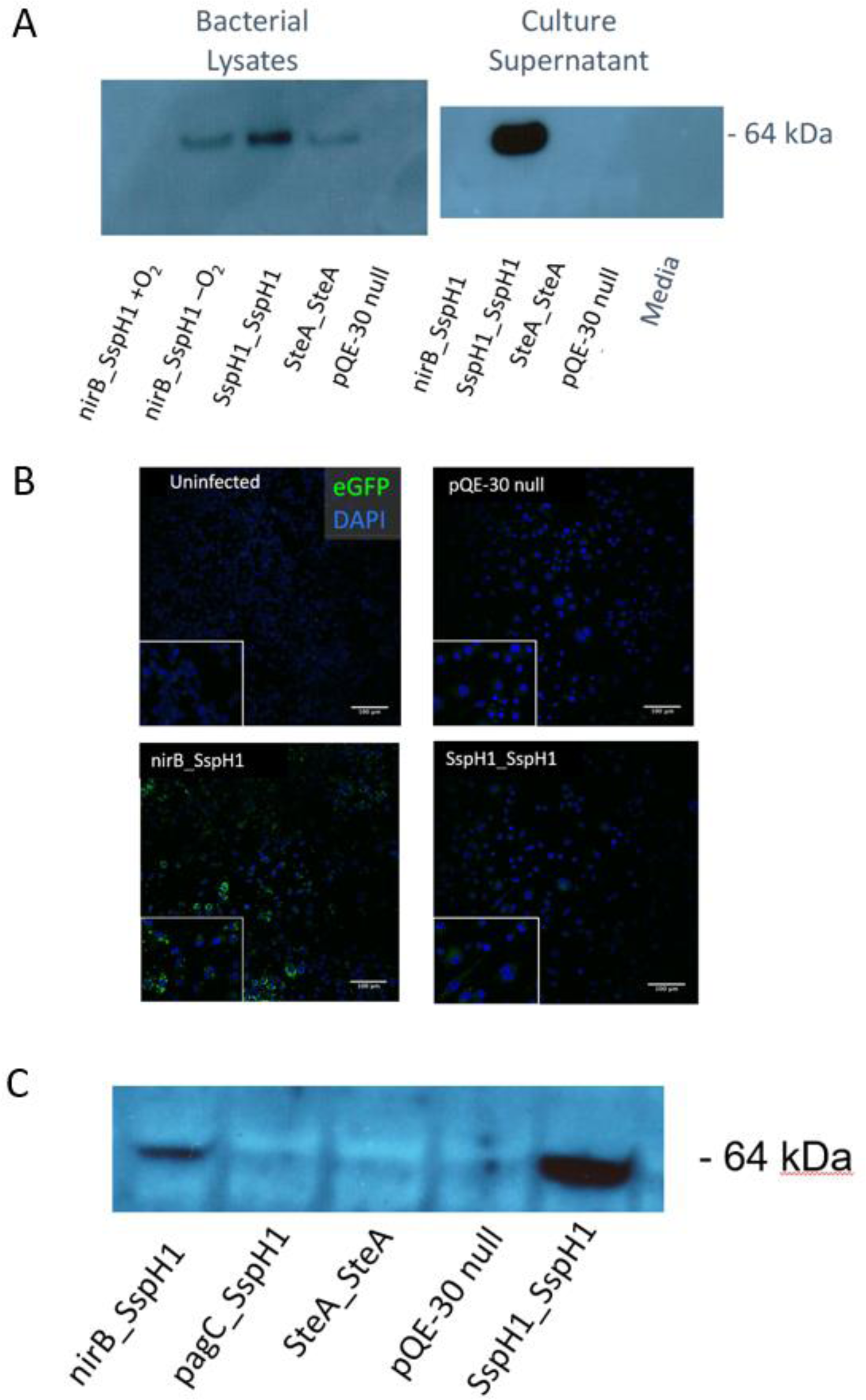
Expression of recombinant cathepsin B. A) The plasmids nirB_SspH1, SspH1_SspH1 and SteA_SteA were transformed into *Salmonella* strain YS1646. Whole bacteria lysates and monomicrobial culture supernatants were examined for the presence of CatB by western blot. B) The mouse macrophage cell line RAW 264.7 cells were infected with transformed YS1646 strains expressing eGFP as a marker for the capacity of promoter-TSSS pairs to support expression of a foreign protein. DAPI nuclear stain is represented in blue and eGFP is shown in green. Scale at 100 μm. C) Mouse macrophage cells line RAW 264.7 cells were infected with selected plasmids from Table 1 and the presence of CatB protein was determined by western blotting.

### Antibody Response to YS1646-vectored Vaccination

None of the groups had detectable anti-CatB antibodies at baseline and the saline control mice remained negative after vaccination. Mice in the PO→PO group also had very low serum CatB-specific antibody levels even after the second vaccination (395.7 ± 48.9: Fig 4A). In contrast, all animals that had received at least 1 dose of rCatB IM had significantly higher IgG titers at 6 weeks (ie: 3 weeks after the second immunization) (Fig 4A). Mice that received nirB_SspH1 PO followed by an IM boost had the highest titers (6766 ± 2128 ng/mL, *P*<.01 vs. control) but these titers were not significantly different from groups that had received either one (EV→IM) or two doses of rCatB (IM→IM) (5898 ± 1951 ng/mL and 6077 ± 4460 ng/mL respectively, both *P*<.05 vs. control). Antibody titers were generally lower in all groups that received the YS1646 strain bearing the SspH1_SspH1 construct (range 333.5 – 3495 ng/mL; *P*<.05, *P*<.01, *P*<.001 vs control: Fig 4B). Neither IgG subtypes nor intestinal IgA levels were measured in the SspH1_SspH1 groups.

**Figure 4.**
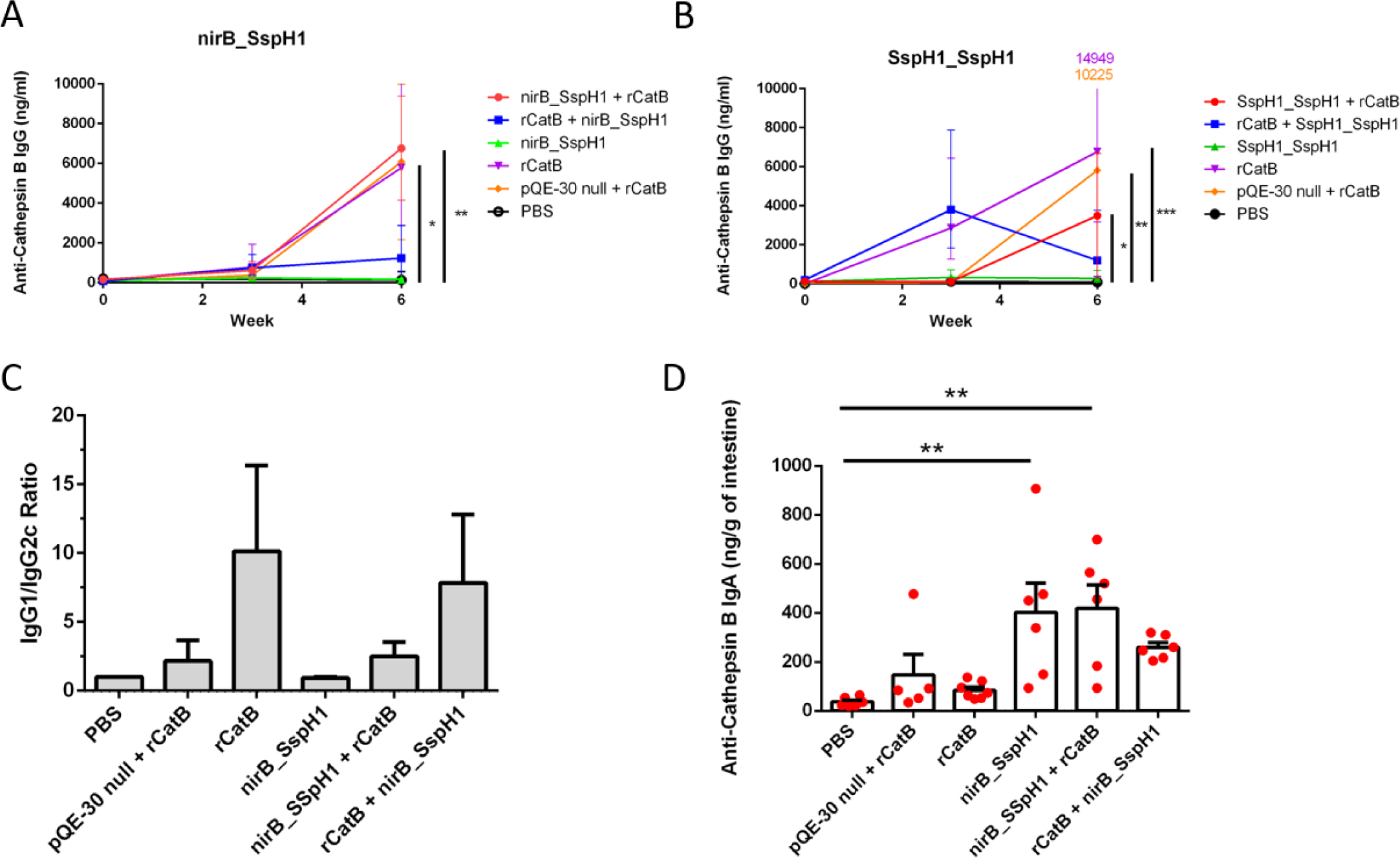
Production of Sm-Cathepsin B specific antibodies prior to challenge. Serum anti-CatB IgG was measured by ELISA at weeks 0, 3 and 6 for groups that received the nirB_SspH1 construct (A) or the SspH1_SspH1 construct (B). These results represent between 8 – 16 animals/group from 2 independent experiments and are reported as the geometric mean with 95% confidence intervals. Significance bars for A and B are to the right of each graph. C) Serum anti-CatB IgG1 and IgG2c were measured by endpoint-dilution ELISA and expressed as the ratio of IgG1/IgG2c. D) Intestinal anti-CatB IgA in intestinal tissue was measured by ELISA and is reported as mean ± standard error of the mean ng/gram. These results represent 5 – 7 animals per group. (\**P<*.05, \*\**P*<.01, \*\**\*P*<.001 compared to the PBS group)

Control mice had no detectable anti-CatB antibodies and were arbitrarily assigned an IgG1/IgG2c ratio of 1. The PO→PO mice had a ratio of 0.9 (Fig 4C). The EV→IM and the PO→IM groups had IgG1/IgG2c ratios of 2.2 and 2.5 respectively while the highest ratios were seen in the IM→IM and IM→PO groups (10.2 and 7.8 respectively).

Intestinal IgA levels in the saline, EV→IM, and IM→IM groups were all low (range 37.0 – 148.0 ng/g of tissue: Fig 4D). Although the data are variable, groups that received at least one dose of nirB_SspH1 YS1646 PO had increased IgA levels compared to the control group that reached statistical significance in the PO→PO group (402.7 ± 119.7 ng/g) and the PO→ IM group (ie: nirB_SspH1 PO then rCatB IM: 419.6 ± 95.3 ng/g, both *P*<.01). The IM→PO group also had higher intestinal IgA titers than controls, but this increase did not reach statistical significance (259.8 ± 19.4 ng/g).

### Cytokine Production in Response to YS1646-vectored Vaccination

There was only modest evidence of CatB-specific cytokine production by antigen re-stimulated splenocytes immediately prior to challenge (4 weeks after the second dose). There were no significant differences in the levels of IL-2, IL-4, IL-10, IL-12p70, IL-13, IL-17 or TNF-α between vaccinated and control groups (S1 Table). Compared to the control group, the levels of IL-5 in splenocyte supernatants were significantly higher in mice that received two doses of rCatB (IM→IM) (475.5 ± 98.5 pg/mL, *P*<.01) and the nirB_SspH1 PO→IM group (364.4 ± 85.2 pg/mL, *P*<.05) whereas the control group was below the limit of detection at 63.1 pg/mL (Fig 5A). Only the PO→IM group had clear evidence of CatB-specific production of IFNγ in response to vaccination (933 ± 237 pg/mL vs. control 216.4 ± 62.5 pg/mL, *P*<.05) (Fig 5B).

**Figure 5.**
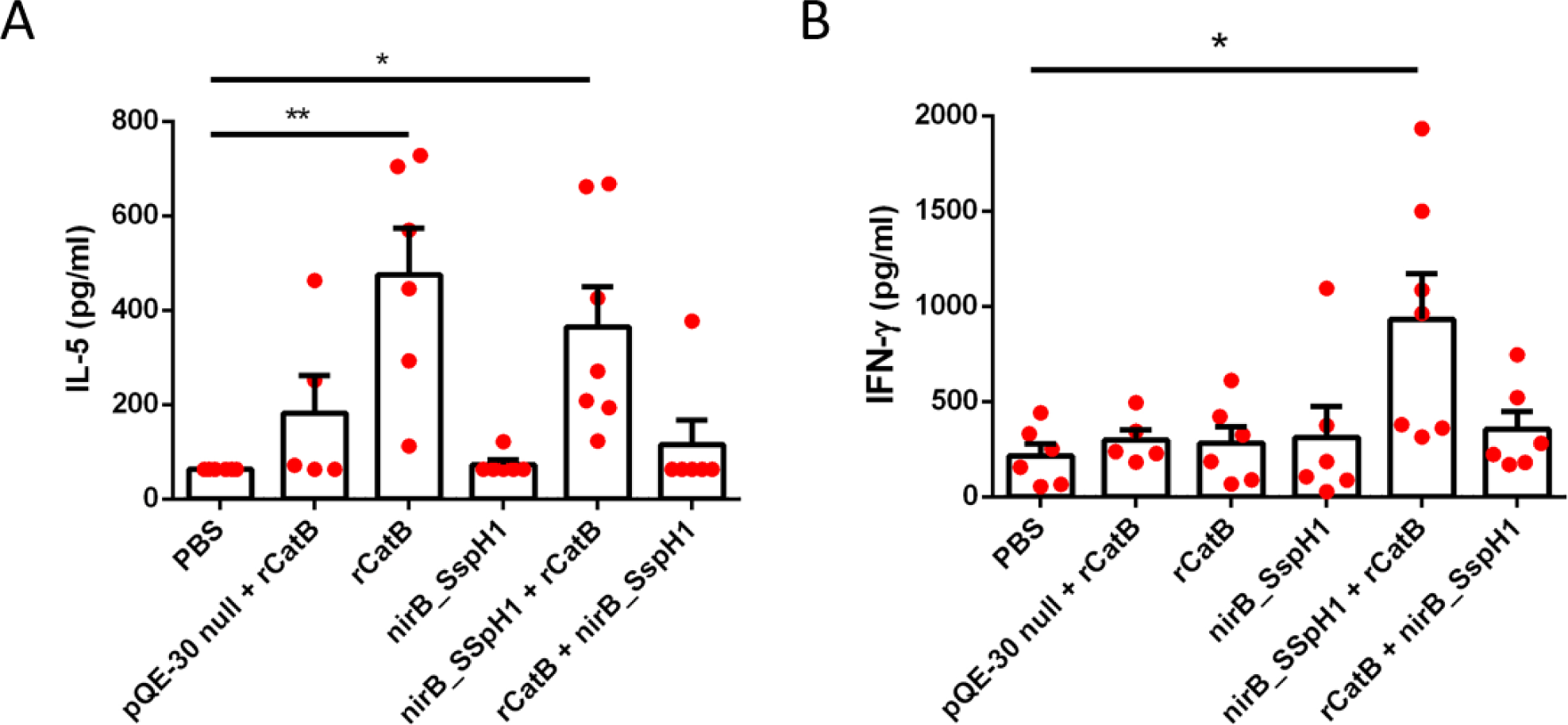
Cytokine production prior to challenge. Supernatant IL-5 (A) and IFN-γ (B) levels after stimulating splenocytes with rCatB for 72 hours were measured by QUANSYS multiplex ELISA. These results represent 5 – 7 animals per group. Results are expressed as the mean ± the standard error of the mean. (\**P<*.05, \*\**P*<.01 compared to the PBS group)

### Protection from *S. mansoni* Challenge from YS1646-vectored Vaccination

At 7 weeks after infection, the mean worm burden in the saline-vaccinated control group was 25.2 ± 4.3. Relatively small reductions in worm burden were observed in the EV→IM (9.4%) and IM→IM groups (20.5%) across all studies. Overall, protection was better with nirB_SspH1_CatB schedules compared to SspH1_SspH1_CatB schedules. In the SspH1_SspH1 animals, reductions in worm numbers were similar to the IM→IM group: 17.2% with oral vaccination alone (PO→PO) and only 17.8% and 24.7% in the PO→IM and IM→PO groups respectively. In contrast, the PO→PO group vaccinated with the nirB_SspH1 YS1646 strain had an 81.7% (*P*<.01) reduction in worm numbers and multi-modality vaccination with this strain achieved 93.1% (*P*<.001) and 81.7% (*P*<.01) reductions in the PO→IM and IM→PO groups respectively. (Fig 6A).

**Figure 6.**
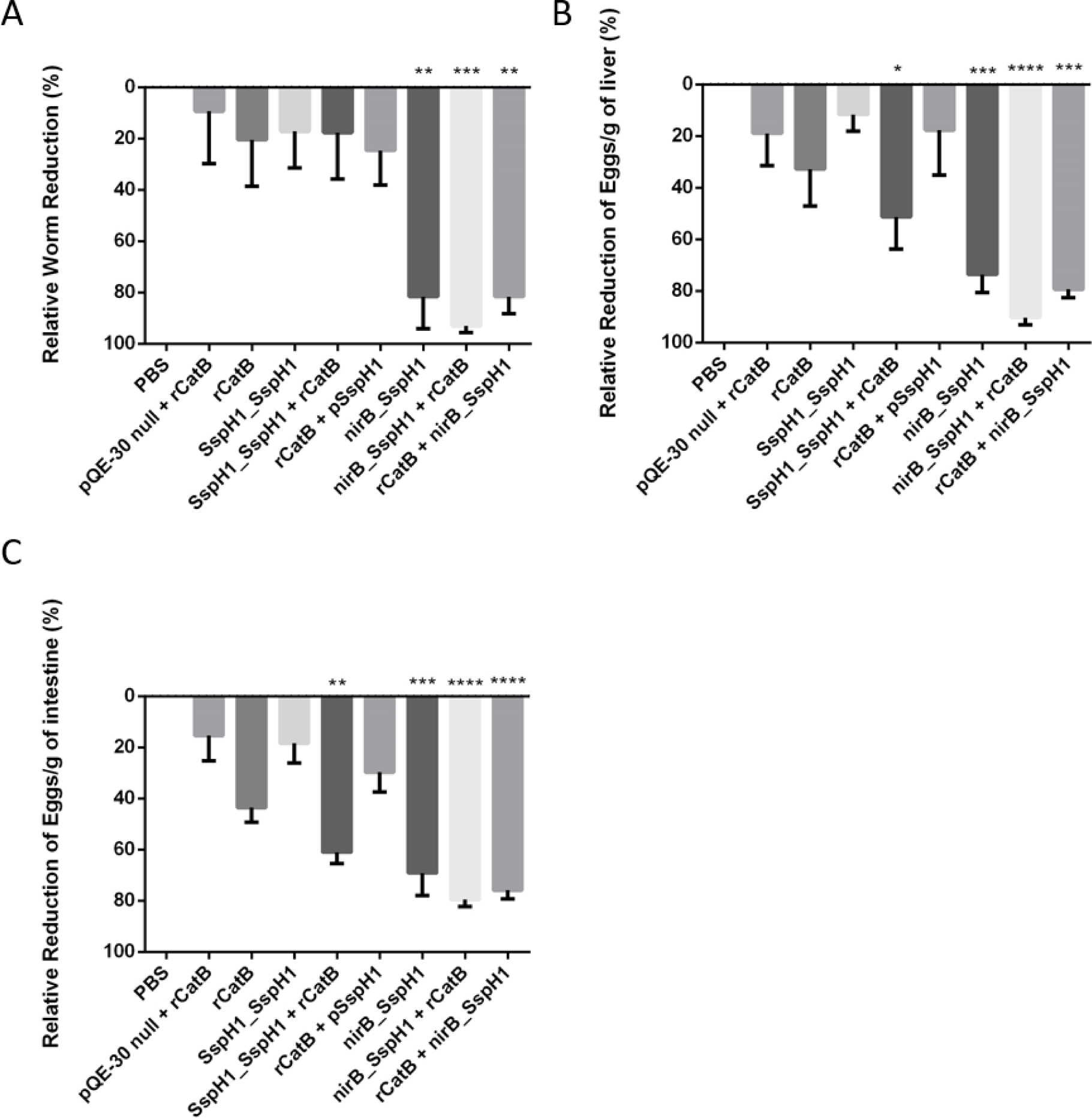
Parasitologic burden. The reduction in worm counts (A) as well as the reduction in egg load per gram of liver (B) or intestine (C) are represented for mice in the PBS, empty vector, PO→PO, IM→IM, and multimodality groups. Worm and egg burdens were determined 7 weeks after cercarial challenge. These results represent between 8 – 16 animals/group from 2 independent experiments. (\**P*<.05, \*\**P*<.01, \*\*\**P*<.001, \*\*\*\**P*<.0001 compared to the PBS group)

Overall, the reductions in hepatic and intestinal egg burden followed a similar pattern to the vaccine-induced changes in worm numbers. The hepatic and intestinal egg burden in the saline-vaccinated control mice ranged from 1,994 – 13,224 eggs/g and 6,548 – 24,401 eggs/g respectively. Reductions in hepatic eggs in the EV→IM and IM→IM groups were modest at 18.9% and 32.7% respectively. Reductions in intestinal eggs followed a similar trend: 15.4% and 43.6% respectively. In the groups that received the SspH1_SspH1 YS1646 strain, PO→PO immunization did not perform any better with 11.6% and 18.3% reductions in hepatic and intestinal egg numbers respectively. Somewhat greater reductions in hepatic and intestinal egg burden were seen in the PO→IM (51.3% and 60.9% respectively) and IM→PO groups (17.7% and 29.8% respectively). These apparent differences in egg burden between the two multi-modality groups did not parallel the reductions in worm numbers or the systemic anti-CatB IgG levels. Groups that received the nirB_SspH1 strain had more consistent and greater reductions in egg burden: the PO→PO group had 73.6% and 69.2% reductions in hepatic and intestinal egg numbers respectively (both *P*<.001). The greatest impact on hepatic and intestinal egg burden was seen in the nirB_SspH1 multi-modality groups: 90.3% (*P*<.0001) and 79.5% (*P*<.0001) respectively in the PO→IM group and 79.4% (*P*<.001) and 75.9% (*P*<.0001) respectively in the IM→PO group (Figs 6B and 6C).

Hepatic granulomas were large and well-formed in the PBS-treated control mice (62.2 ± 6.1 x10^3^ μm^2^) and essentially all of the eggs in these granulomas had a normal appearance. The EV→IM and IM→IM groups had slightly smaller granulomas (52.0 ± 6.9 x10^3^ μm^2^ and 52.8 ± 10.4 x10^3^ μm^2^ respectively) with modest numbers of abnormal-appearing eggs (ie: loss of internal structure, crenellated edge) (Table 2) but these differences did not reach statistical significance. Groups that received the SspH1_SspH1 strain had granuloma sizes ranging from 47.3–55.0 x10^3^ μm^2^ with 30.5% of the eggs appearing abnormal in the PO→IM and 28.6% IM→PO groups (both *P*<.05). In the groups that received the nirB_SspH1 strain, both the purely oral (PO→PO) and multi-modality strategies (PO→IM and IM→PO) resulted in even smaller granulomas (32.9 ± 2.0 μm^2^, 34.7 ± 3.4 x10^3^ μm^2^ and 39.2 ± 3.7 x10^3^ μm^2^: *P*<.01, *P*<.01 and *P*<.05 respectively). The large majority of the eggs in these granulomas had disrupted morphology (75.9 ± 7.6%, 79.4 ± 4.2% and 71.9 ± 6.0% respectively: all *P*<.0001). Overall, the greatest and most consistent reductions in both adult worm numbers and egg burdens in hepatic and intestinal tissues were seen in the animals that received oral dosing with the YS1646 bearing the nirB_SspH1_CatB construct followed 3 weeks later by IM rCatB.

**Table 2.** Granuloma size and egg morphology.

| Group | Granuloma size ($\times 10^3 \mu\text{m}^2$)<br>± SEM | Abnormal egg morphology<br>(%) ± SEM |
| --- | --- | --- |
| PBS | $62.2 \pm 6.1$ | 0 |
| pQE-30 null + rCatB | $52.0 \pm 6.9$ | $18.9 \pm 3.9$ |
| rCatB | $52.8 \pm 10.4$ | $12.6 \pm 5.1$ |
| SspH1_SspH1 | $55.0 \pm 8.5$ | $25.0 \pm 6.2$ |
| SspH1_SspH1 + rCatB | $47.3 \pm 4.4$ | $30.5 \pm 7.7^*$ |
| rCatB + SspH1_SspH1 | $49.8 \pm 14.3$ | $28.6 \pm 6.8^*$ |
| nirB_SspH1 | $32.9 \pm 2.0^{**}$ | $75.9 \pm 7.6^{****}$ |
| nirB_SspH1 + rCatB | $34.7 \pm 3.4^{**}$ | $79.4 \pm 4.2^{****}$ |
| rCatB + nirB_SspH1 | $39.2 \pm 3.7^*$ | $71.9 \pm 6.0^{****}$ |

Liver granuloma area ( $\times 10^3 \mu\text{m}^2$ ) and egg morphology (ie: loss of internal structures, shrinkage, crenelated periphery) were assessed. SEM represents the standard error of the mean. (\* $P<.05$ , \*\* $P<.01$ , \*\*\*\* $P<.0001$ compared to the PBS group)

## Discussion

In the mid-1990s, the Tropical Diseases Research (TDR) committee of the World Health Organization (WHO) launched the search for a *S. mansoni* vaccine candidate capable of providing ≥40% protection [9]. This initiative targeted reduced worm numbers as well as reductions in egg burden in both the liver and the intestinal tissues. *S. mansoni* female worms can produce hundreds of eggs per day [33]. While the majority are excreted in the feces, some are trapped in host tissues where they cause most of the pathology associated with chronic infection [34]. Eggs trapped in the liver typically induce a vigorous granulomatous response that can lead to fibrosis, cirrhosis and death while egg-induced granulomas in the intestine cause local lesions that contribute to colonic polyp formation [35]. Reducing the hepatic egg burden would therefore be predicted to decrease *S. mansoni*-associated morbidity and mortality while reducing the intestinal egg burden would likely decrease transmission.

Our group has previously described the protective efficacy of CatB-based vaccines delivered IM with adjuvants. Using CpG dinucleotides to promote a Th1-type response, vaccination resulted in a 59% reduction in worm burden after challenge with 56% and 54% decreases in hepatic and intestinal egg burden respectively compared to adjuvant-alone control animals [12]. Parasitologic outcomes were slightly better in the same challenge model when the oil-in-water adjuvant Montanide ISA 720 VG was used to improve the antibody response: 56-62% reductions in worm numbers and the egg burden in tissues [13]. These results were well above the 40% threshold suggested by the TDR/WHO and provided proof-of-concept for CatB as a promising target antigen. Based on this success, we expanded our vaccine discovery program to explore alternate strategies and potentially more powerful delivery systems. The availability of the highly attenuated *Salmonella enterica* Typhimurium strain YS1646 that had been used in a phase 1 clinical cancer trial at doses up to 3×10^8^ IV was attractive for many reasons. Although *S. enterica* species replicate in a membrane-bound host cell compartment or vacuole [36], foreign protein antigens can be efficiently exported from the vacuole into the cytoplasm using the organism’s T3SS. Like all *Salmonella enterica* species, YS1646 has two distinct T3SS located in *Salmonella* pathogenicity islands 1 and 2 (SPI-I and SPI-II) [37] that are active at different phases of infection [38]. The SPI-I T3SS translocates proteins upon first contact of the bacterium with epithelium cells through to the stage of early cell invasion while SPI-II expression is induced once the bacterium has been phagocytosed [39]. These T3SS have been used by many groups to deliver heterologous antigens in *Salmonella*-based vaccine development programs [22, 40].

In this study, we report the protective efficacy of CatB delivered by the attenuated strain YS1646 of *Salmonella enterica* serovar Typhimurium in a heterologous prime-boost vaccination regimen. Compared to infected controls, vaccination with CatB IM followed by YS1646 bearing the nirB_SspH1 strain resulted in an 93.1% reduction in worm numbers and 90.3% and 79.5% reductions in hepatic and intestinal egg burdens respectively. These results not only surpass the WHO’s criterion for an effective *S. mansoni* vaccine by a considerable margin, they are a marked improvement on our own work using CatB delivered IM with adjuvants and are among the best results ever reported in similar murine models [12, 13]. For example, in the pre-clinical development of two candidate vaccines that subsequently entered clinical trials [41, 42], IM administration of the fatty acid binding protein Sm-14 with the adjuvant GLA-SE led to a 67% reduction in worm burden in mice [10] while IM vaccination with the tegumental protein TSP-2 with either Freund’s adjuvant or alum/CpG reduced worm numbers by 57% and 25% and hepatic egg burden by 64% and 27% respectively [43, 44]. Another vaccine candidate targeting the tegumental protein Sm-p80 that is advancing towards clinical testing achieved 70 and 75% reductions in adult worm numbers and hepatic egg burden respectively when given IM with the oligodeoxynucleotide (ODN) adjuvant 10104 [45]. It is noteworthy that these other vaccine candidates were all administered IM. Although this route would be expected to generate high systemic antibody titers, particularly with the use of adjuvants, it is unlikely that any would elicit a local, mucosal response like the multimodality approach taken in our studies.

To what extent the surprising reductions in worm and egg burdens that we observed with the YS1646 can be attributed to the systemic or the local antibody response is currently unknown although it is likely that both contributed to the success of the combined schedules (ie: IM→PO and PO→IM). Oral administration of *Salmonella*-vectored vaccines clearly leads to higher mucosal IgA responses than IM dosing [46] and the protective potential of IgA antibodies has been demonstrated in schistosomiasis [47]. The importance of the local response is strongly suggested by the fact that PO dosing alone with YS1646 bearing the nirB_SspH1_CatB construct still provided substantial protection (81.7% and 73.6%/69.2% for worms and hepatic/intestinal eggs) despite the almost complete absence of a detectable systemic response (Fig 4A). Indeed, IgA titers were readily detectable in the intestinal tissues of mice receiving the nirB_SspH1 YS1646 vaccine PO →PO and in mice the received PO→IM dosing (402.7 ng/g and 419.6 ng/g respectively) (Fig 4D). On the other hand, the importance of IgG antibodies in the protection against schistosomiasis has been reported by many groups [48, 49]. Administered IM, rCatB alone consistently elicited high systemic antibody responses and provided a modest level of protection without any measurable mucosal response. Chen and colleagues have also used YS1646 as a vector to test single- and multi-modality approaches for a bivalent vaccine candidate (Sj23LHD-GST) targeting *S. japonicum* in a similar murine model [29]. Although some authors have promoted so-called ‘prime-pull’ strategies to optimize mucosal responses (ie: ‘prime’ in the periphery then ‘pull’ to the target mucosa) [50], it is interesting that both the Chen group and our own findings suggest that PO→IM dosing may be the optimal strategy. In the *S. japonicum* model targeting the long hydrophobic domain of the surface exposed membrane protein Sj23LHD and a host-parasite interface enzyme (glutathione S-transferase or GST), the PO→IM vaccination schedule led to important reductions in both worm numbers (51.4%) and liver egg burden (62.6%) [29].

In addition to the substantial overall reductions in worm numbers and egg burden in our animals that received multimodality vaccination, there were additional suggestions of benefit in terms of both hepatic granuloma size and possible reduced egg fitness (Table 2). The size of liver granulomas is determined largely by a Th2-deviated immune response driven by soluble egg antigens (SEA) [51]. Prior work with CatB vaccination suggests that IM delivery of this antigen alone tends to elicit a Th2-biased response that can be shifted towards a more balanced Th1/Th2 response by CpG or Montanide [12, 13, 52]. The reduction in the anti-CatB IgG1/IgG2c ratio between the IM→IM only and multimodality groups (IM→PO, PO→IM) supports the possibility that combined recombinant CatB with YS1646 bearing CatB can induce a more ‘balanced’ pattern of immunity to this antigen and, at least in a limited sense, that the YS1646 is acting as a Th1-type adjuvant (Fig 4C). Although no adjuvants were included in the current study, the YS1646 vector might reasonably be considered ‘auto-adjuvanted’ by the presence of LPS, even in an attenuated form, and flagellin which can act as TLR-4 and TLR-5 agonists respectively. It was still surprising however, that the average hepatic granuloma size was significantly smaller in our multi-modality groups than in the IM alone group since no CatB is produced by the eggs (Table 2). This observation raises the interesting possibility that the YS1646-based vaccination protocol may be able to influence the *overall* pattern of immunity to *S. mansoni* and/or reduce the fitness of the eggs produced (as suggested by the abnormal egg morphology observed). Such effects could significantly extend the value of the combined PO→IM vaccination strategy, i.e.: more durable impact, reduced transmission, etc. Furthermore, prior work with IM vaccination with CatB alone revealed a Th2-type pattern of cytokine response in splenocytes (eg: IL-4, IL-5, and IL-13) [52]. In the current work we observed increases in both IFNγ and IL-5 in the multimodality PO→IM group (Fig 5), suggesting that YS1646 vaccination can induce more balanced Th1-Th2 immune response. Finally, this study did not consider the possible role of other immune mechanisms in controlling *S. mansoni* infection after YS1646 infection and we have previously shown that CD4^+^ T cells and anti-schistosomula antibody-dependent cellular cytotoxicity (ADCC) contribute to protection after CatB immunization (± adjuvants) [53]. Studies are underway to examine these possibilities with the multi-modality YS1646-based vaccination protocols. It is also intriguing that the apparent efficacy of either one or two IM doses of rCatB differed considerably between the EV→IM and IM→IM groups with the latter schedule eliciting significantly greater protection for all parasitologic outcomes despite the fact that these groups had similar levels of serum anti-CatB IgG at the time of challenge (Fig 4). Future studies will address whether or not there are qualitative differences in the antibodies induced (ie: avidity, isotype, competence to mediate ADCC) and/or differences in other immune effectors (ie: CD4^+^ or CD8^+^ T cells).

Although the findings presented herein are promising, this work has several limitations. First, immune protection is likely to be relatively narrow when only a single schistosome antigen is targeted. In the long term, this limitation could be easily overcome by adding one or more of the many *S. mansoni* target antigens that have shown promise in pre-clinical and/or clinical development (eg: GST, Sm23, Sm-p80, etc) to generate a ‘cocktail’ vaccine. In this context, an attenuated *Salmonella* vector like YS1646 might be ideal because of its high ‘carrying capacity’ for foreign genes [54]. Second, our current findings are based on plasmid-mediated expression and pQE30 contains a mobile ampicillin resistance gene that would obviously be inappropriate for use in humans [55]. Although chromosomal integration of our nirB_SspH1_CatB gene is an obvious mitigation strategy, expression of the CatB antigen from a single or even multiple copies of an integrated gene would likely be lower than plasmid-driven expression. Finally, the degree to which a vaccination schedule based on the YS1646 vector would be accepted by regulators is currently unknown. Attenuated *Salmonellae* have a good safety track-record in vaccination: eg: the Ty21a *S. typhi* vaccine and a wide range of candidate vaccines [54] despite their ability to colonize/persist for short periods of time [56]. Although the total clinical exposure to YS1646 to date is limited (25 subjects with advanced cancer in a phase 1 anti-cancer trial), the available data are reassuring since up to 3×10^8^ bacteria could be delivered intravenously in these vulnerable subjects without causing serious side effects [16].

In summary, this work demonstrates that a YS1646-based, multimodality, prime-boost immunization schedule can provide nearly complete protection against *S. mansoni* in a well-established murine model. The protection achieved against a range of parasitologic outcomes was the highest reported to date for any vaccine. These observations strongly support the continued development of this candidate vaccine for prophylactic and possibly even therapeutic use in the many hundreds of millions of people in low- and middle-income countries at risk of or already infected by *S. mansoni*.

## Acknowledgements

We thank Annie Beauchamp for assistance with all animal work, Lydia Labrie for assistance with sample collection and processing, and Louis Cyr for assistance with the histopathology staining. We also would like to thank Dr. Margaret Mentink-Kane and Kenia V. Benitez from the Biomedical Research Institute (Rockville, MD) for supplying us with infected *Biomphalaria* snails.

## Authors’ Contributions

ASH was involved in all aspects of the study including study design, performing experiments, data analysis and preparation of the manuscript. NHZ designed, constructed and validated the plasmids. DJP assisted in the design of the snail housing facility and the infection model. BJW and MN supervised all parts of the study and prepared the manuscript.

## Supporting information

**S1 Table.**
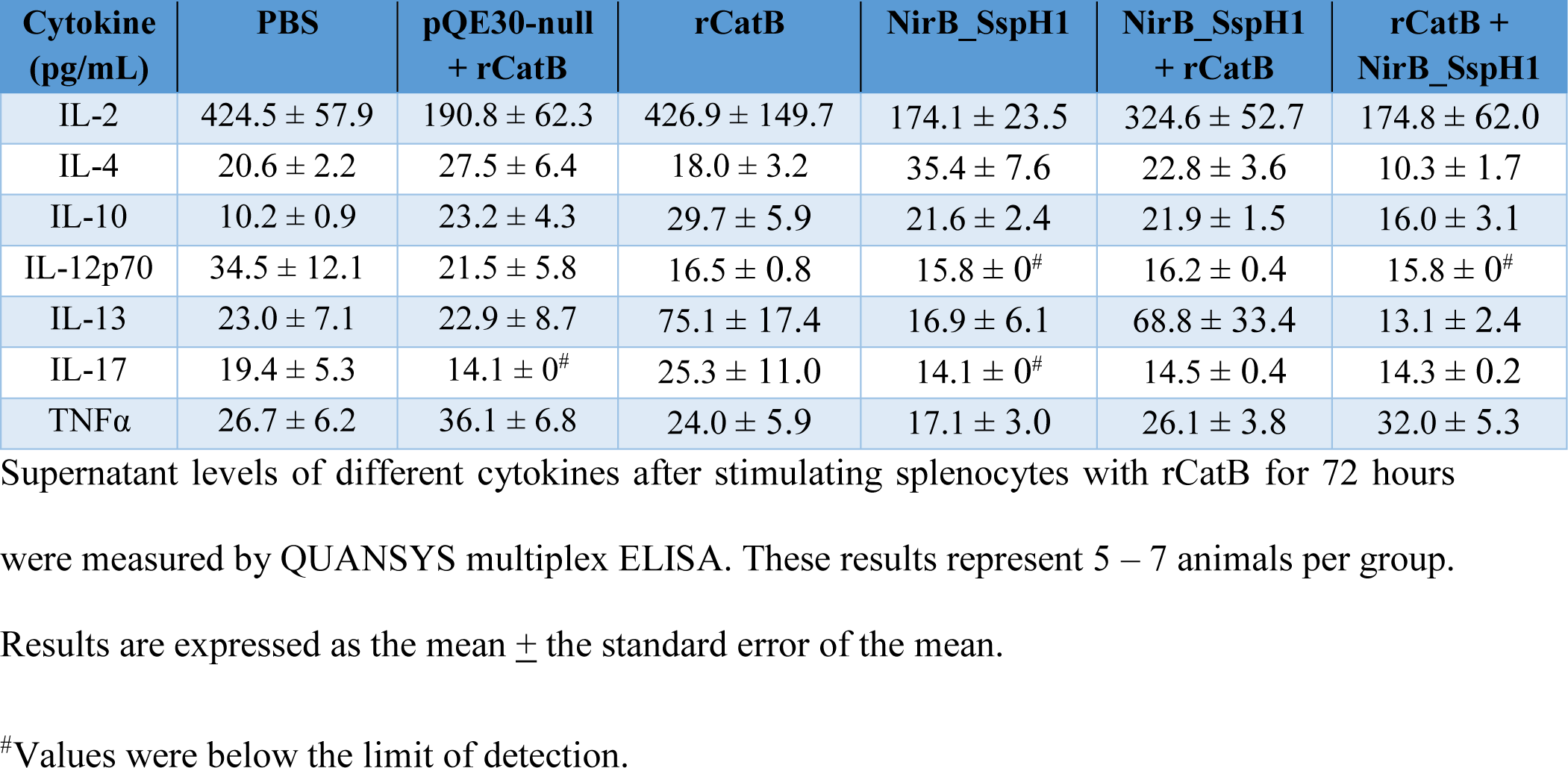
Other cytokine production prior to challenge.

| Cytokine (pg/mL) | PBS | pQE30-null + rCatB | rCatB | NirB_SspH1 | NirB_SspH1 + rCatB | rCatB + NirB_SspH1 |
| --- | --- | --- | --- | --- | --- | --- |
| IL-2 | 424.5 ± 57.9 | 190.8 ± 62.3 | 426.9 ± 149.7 | 174.1 ± 23.5 | 324.6 ± 52.7 | 174.8 ± 62.0 |
| IL-4 | 20.6 ± 2.2 | 27.5 ± 6.4 | 18.0 ± 3.2 | 35.4 ± 7.6 | 22.8 ± 3.6 | 10.3 ± 1.7 |
| IL-10 | 10.2 ± 0.9 | 23.2 ± 4.3 | 29.7 ± 5.9 | 21.6 ± 2.4 | 21.9 ± 1.5 | 16.0 ± 3.1 |
| IL-12p70 | 34.5 ± 12.1 | 21.5 ± 5.8 | 16.5 ± 0.8 | 15.8 ± 0<sup>#</sup> | 16.2 ± 0.4 | 15.8 ± 0<sup>#</sup> |
| IL-13 | 23.0 ± 7.1 | 22.9 ± 8.7 | 75.1 ± 17.4 | 16.9 ± 6.1 | 68.8 ± 33.4 | 13.1 ± 2.4 |
| IL-17 | 19.4 ± 5.3 | 14.1 ± 0<sup>#</sup> | 25.3 ± 11.0 | 14.1 ± 0<sup>#</sup> | 14.5 ± 0.4 | 14.3 ± 0.2 |
| TNFα | 26.7 ± 6.2 | 36.1 ± 6.8 | 24.0 ± 5.9 | 17.1 ± 3.0 | 26.1 ± 3.8 | 32.0 ± 5.3 |

Supernatant levels of different cytokines after stimulating splenocytes with rCatB for 72 hours were measured by QUANSYS multiplex ELISA. These results represent 5 – 7 animals per group. Results are expressed as the mean ± the standard error of the mean.
<sup>#</sup>Values were below the limit of detection.

**S2 Table.**
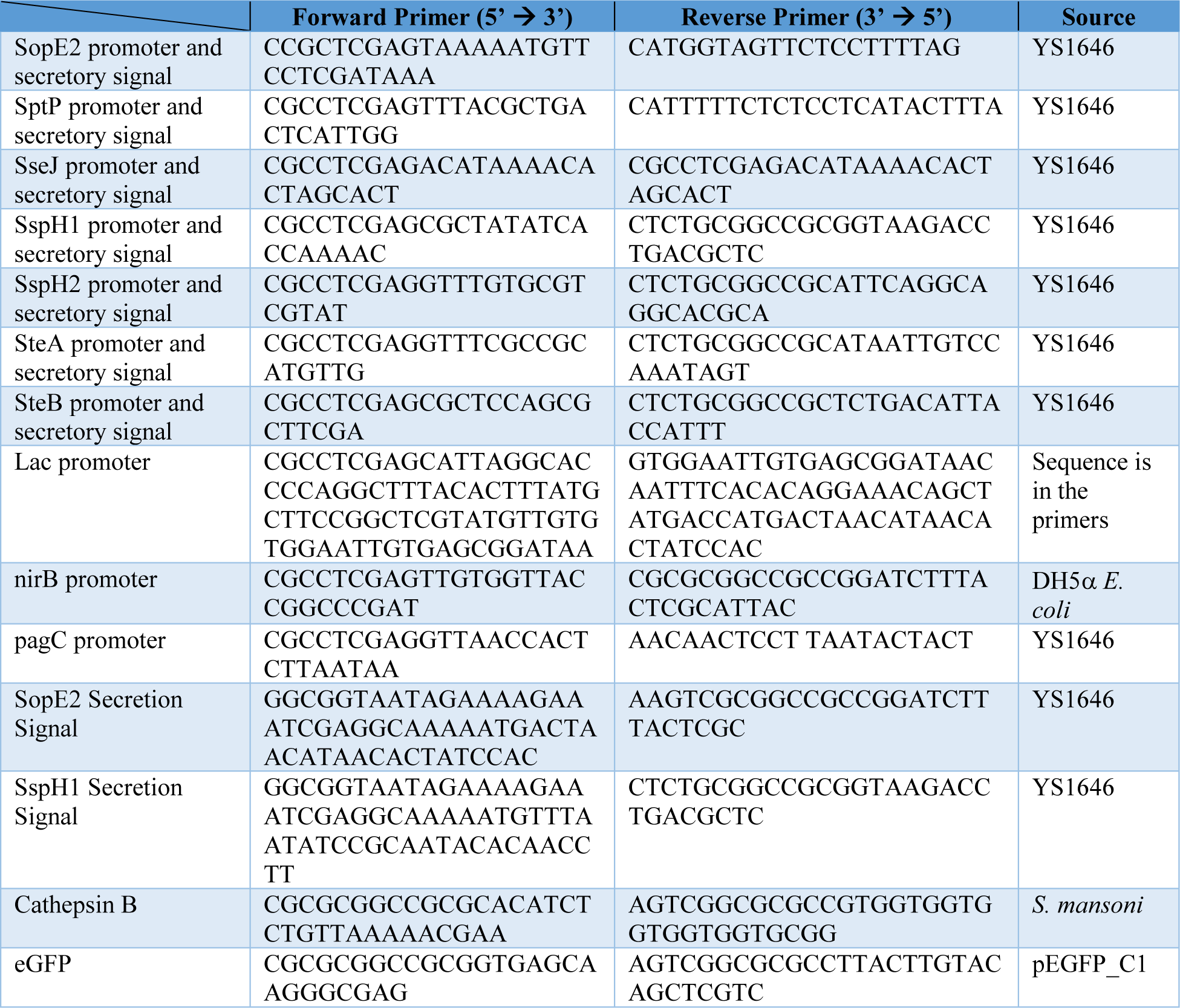
Primers used in construct design.

